# The relationship between misfolding avoidance hypothesis and protein evolutionary rates in the light of empirical evidence

**DOI:** 10.1101/736280

**Authors:** Dinara R. Usmanova, Germán Plata, Dennis Vitkup

**Author notes:** These authors contributed equally (Dinara R. Usmanova, Germán Plata). Correspondence to DV.

## Abstract

For more than a decade the misfolding avoidance hypothesis (MAH) and related theories have dominated evolutionary discussions aimed at explaining the variance of molecular clock across cellular proteins. In this study we use various experimental data to further investigate the consistency of the MAH predictions with empirical evidence. We also critically discuss experimental results that motivated the MAH development and that are often viewed as evidence of its major contribution to constraining protein evolution. We demonstrate, in *Escherichia coli* and *Homo sapiens*, the lack of a substantial negative correlation between protein evolutionary rates and Gibbs free energies of unfolding, a direct measure of protein stability. We then analyze multiple new genome-scale datasets describing protein aggregation and interaction propensities, which are likely optimized in evolution to alleviate deleterious effects associated with toxic protein misfolding and misinteractions. Our results demonstrate that the propensity of proteins to aggregate, the fraction of charged amino acids, and protein stickiness do correlate with protein abundances. Nevertheless, across multiple organisms and datasets we do not observe substantial correlations between proteins aggregation- and stability-related properties and evolutionary rates. Therefore, diverse empirical data support the conclusion that the MAH and similar hypotheses are unlikely to play a major role in mediating a strong negative correlation between protein expression and molecular clock, and thus in explaining the variability of evolutionary rates across cellular proteins.

**Significance statement:** Evolutionary rates vary substantially across cellular proteins. Understanding the nature of molecular clock and its variability across proteins is a foundational question in molecular evolution. The popular and currently dominant theory to explain the molecular clock variability is the misfolding avoidance hypothesis (MAH). The role of the MAH is currently under active debate. In the manuscript we discuss how to appropriately test the MAH based on available empirical data, and then rigorously test the hypothesis using more than a dozen of new genome-wide datasets of protein stability and aggregation propensities. Our results suggest that the MAH is unlikely to play a major role in explaining the variability of molecular clock across proteins.

## Introduction

Protein evolutionary rates vary by orders of magnitude, but the mechanisms underlying this variability are currently unknown (Koonin 2012). Although protein expression was shown to be the strongest predictor of protein evolutionary rates across species (Pal, et al. 2001; Pal, et al. 2006), the causes of the anticorrelation between expression and protein evolutionary rate are not understood (Zhang and Yang 2015). The popular misfolding avoidance hypothesis (MAH) posits that the sequences of highly abundant proteins evolve slowly *primarily* due to increased selection against misfolded protein toxicity (Drummond, et al. 2005; Drummond and Wilke 2008; Yang, et al. 2010). The recent availability of genome-wide experimental data on protein stability has reinvigorated the debate about the model of protein evolution based on the MAH (Plata and Vitkup 2018; Razban 2019).

We read with interest the recent manuscript “Protein melting temperature cannot fully assess whether protein folding free energy underlies the universal abundance-evolutionary rate correlation seen in proteins” by Rostam Razban (Razban 2019). This paper is related to our previous analyses, i.e. Plata *et al.* (2010) (Plata, et al. 2010) and especially Plata and Vitkup (2018) (Plata and Vitkup 2018). Razban’s manuscript discusses our results showing a lack of empirical support for the MAH based on the genome-wide protein melting temperature (*T*_*m*_) data obtained by Leuenberger *et al.* (Leuenberger, et al. 2017). To avoid potential misunderstanding in the field, in this paper we address inaccuracies in the characterization of our previous work, comment more broadly on the proper usage of experimental data to test the MAH, and then further test the hypothesis using multiple new empirical datasets.

The MAH can be tested by investigating the two key predictions of the hypothesis: (i) protein abundance positively correlates with protein stability (Drummond and Wilke 2008; Zhang and Yang 2015), and (ii) protein stability substantially affects the variation of evolutionary rates across cellular proteins. As we demonstrated previously, a careful re-analysis of the proteome-wide *T*_*m*_ measurements obtained by Leuenberger *et al.* (Leuenberger, et al. 2017) shows no support for the MAH in *E. coli* and three other investigated species (Plata and Vitkup 2018). Razban’s manuscript states that our analysis provides support for the MAH in *E. coli* due to a correlation between protein abundances and melting temperatures. This claim is not correct and, we believe, exemplifies a common and unfortunate confusion. As we specifically discussed (Plata, et al. 2010; Plata and Vitkup 2018), the MAH cannot be validated simply by demonstrating a weak correlation between protein abundance and stability, i.e. the relationship (i) above. The MAH, at its core, is not only about the stability of highly expressed proteins, but also about a major effect of protein stability on the level of sequence constraints across cellular proteins. Thus, it is essential to investigate whether protein stability accounts for any substantial fraction of the variance of evolutionary rates across proteins.

In this study we analyze available Gibbs unfolding free energies (Δ*G*) for *E. coli* and *H. sapiens* proteins (Kumar, et al. 2006), and several proteome-wide datasets characterizing protein aggregation and misinteraction propensities in several species (Levy, et al. 2012; Savitski, et al. 2014; Becher, et al. 2018; Mateus, et al. 2018; Volkening, et al. 2019). Our analyses, based on either the direct correlation between the rate of protein evolution and protein stability, or the partial correlation between the rate of evolution and protein or mRNA abundance, while controlling for stability, show no support for a major role of the MAH in any considered organism.

## Results

The central message of Razban’s analysis is that the absence of the expected relationship between *T*_*m*_ and protein abundance may be due to an imperfect correlation between measured *T*_*m*_ and Δ*G*, based on the error model Razban constructed using the *E. coli* dataset from Leuenberger *et al.* (Leuenberger, et al. 2017; Razban 2019). To address the issue of the correlation between protein melting temperature and stability, we analyzed available data describing protein unfolding Gibbs free energies, Δ*G*. Empirical Δ*G* values have now been obtained for a substantial number of proteins in *E. coli* and *H. sapiens*, and are available in the ProTherm database (Kumar, et al. 2006). This analysis revealed that Razban’s model contradicts a strong empirical correlation for *E. coli* proteins between *T*_*m*_ measurements from Leuenberger *et al.* (Leuenberger, et al. 2017) and Δ*G* values in ProTherm (Pearson’s r=0.69, p-value=0.005; Spearman’s r=0.62, p-value=0.01).

Importantly, a direct measure of protein stability, Δ*G*, allows to test the MAH regardless of an imperfect correlation between *T*_*m*_ and Δ*G*. For the subsets of proteins with available Δ*G* measurements we were able to robustly reproduce a significant anticorrelation between evolutionary rates and mRNA abundances (**Figure 1**, blue; **Supplementary Table 1**; Spearman’s *r* = −0.56, *p* = 2 ∙ 10^−3^, for *E. coli*; *r* = −0.54, *p* = 2 ∙ 10^−4^, for *H. sapiens*). Thus, the mechanisms that make highly expressed proteins evolve more slowly are likely to be reflected in the properties of these proteins. But contrary to the MAH predictions, we did not observe, for *E. coli* and *H. sapiens* proteins, a negative correlation between Δ*G* and evolutionary rates (**Figure 1**, red); we also did not observe positive correlations between Δ*G* and either protein or mRNA abundances (**Figure 1**, light and dark grey, respectively). When we restricted the Δ*G* analysis to include only protein monomers with two-state reversible (un)folding, the relationship between protein stability and abundance in *E. coli* became marginally significant, but in the direction opposite to the one predicted by the MAH (Spearman’s *r* = −0.39, *p* = 0.06; **Supplementary Figure 1**). This pattern may be due to well-known effects associated with the activity-stability tradeoff, i.e. protein functional optimization which often leads to lower protein stability (Wang, et al. 2002; Tokuriki, et al. 2008; Knies, et al. 2017). In summary, in agreement with the conclusions based on *T*_*m*_ measurements from Leuenberger *et al.* (Plata and Vitkup 2018), the empirical Δ*G* data also do not provide any support for a major role of the MAH in explaining the variation of evolutionary rates across proteins.

**Figure 1.**
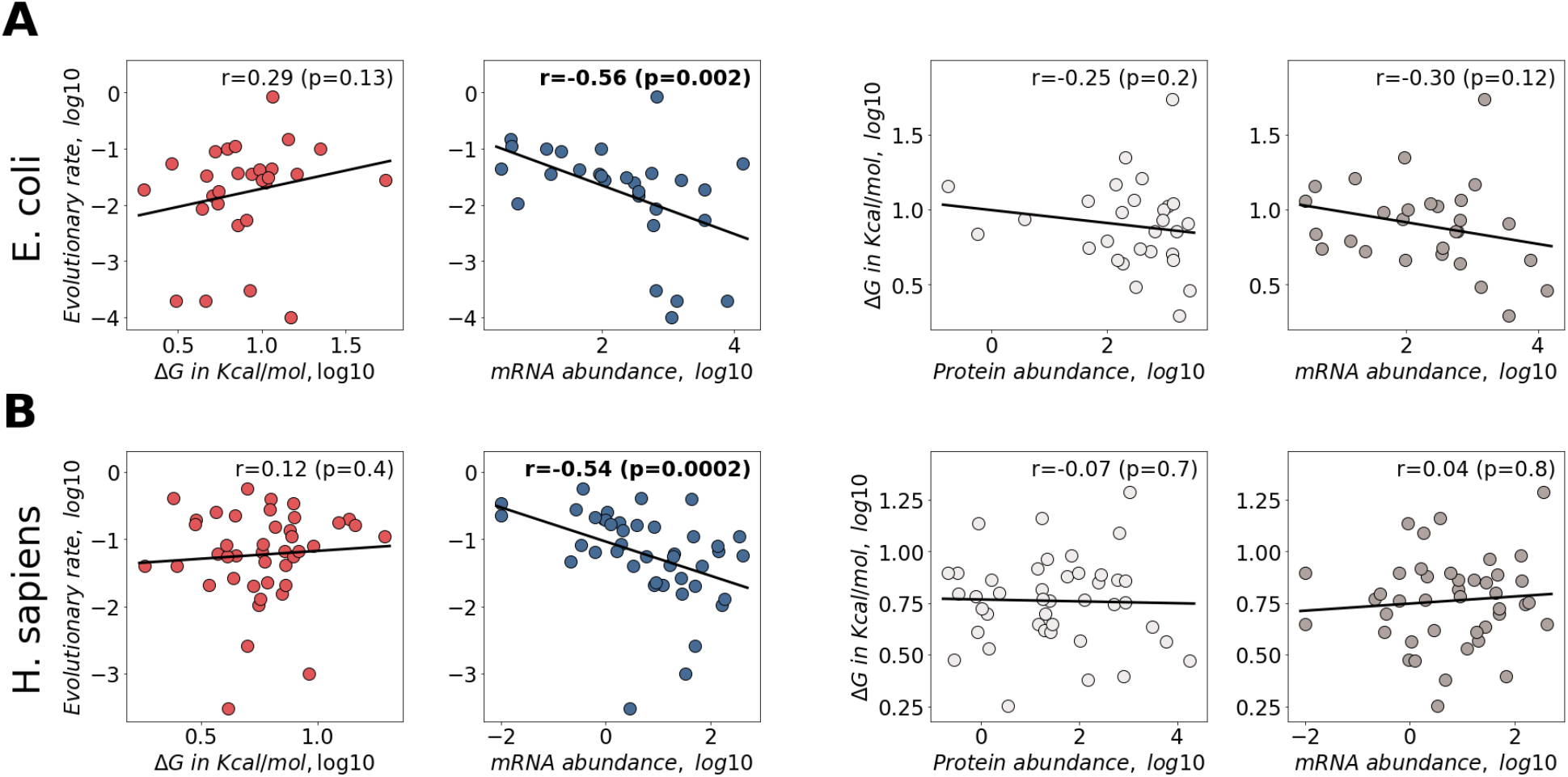
Correlations between experimentally measured Δ*G* values, protein abundances, mRNA abundances, and protein evolutionary rates. **A.** *E. coli* (n=28) and **B.***H. sapiens* (n=42). The correlations between evolutionary rates and unfolding Gibbs free energies, Δ*G*, are shown in the first figure column (red). The correlations between protein evolutionary rates and mRNA abundances are shown in the second column (blue). The correlations between Δ*G* and protein abundances are shown in the third column (light grey), and the correlations between Δ*G* and mRNA abundances are shown in the fourth column (dark grey). Solid lines represent the least square regressions fitted to the data. Spearman’s correlation coefficients and corresponding p-values are shown, significant correlations are highlighted in bold.

In our view, the main caveat with Leuenberger *et al.* dataset, in the context of testing the MAH, is not a poor correlation between *T*_*m*_ and Δ*G*. We previously demonstrated a substantial correlation (Pearson’s *r* = 0.75, *p* < 10^−20^; Spearman’s *r* = 0.64, *p* < 10^−20^) between these two characteristics of protein stability for measurements performed by the same research group (Plata and Vitkup 2018); and as described above, we now also confirmed this correlation specifically for *T*_*m*_ measurements from Leuenberger *et al*. Instead, intrinsic protein stability may simply not serve as a good proxy for protein aggregation propensity, which is likely to mediate misfolding toxicity. The protein melting temperatures obtained by Leuenberger *et al.* (Leuenberger, et al. 2017) are based on data from limited proteolysis (LiP), which increases due to local protein unfolding triggered by higher temperatures; below, we use the term 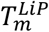 for these melting temperature measurements. An alternative method, developed by Savitski *et al.* (Savitski, et al. 2014), uses protein aggregation (Agg) as a proxy of unfolding. This method estimates melting temperatures by quantifying protein concentrations in soluble cellular fractions as a function of increasing temperature; we use the term 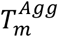 for these melting temperature measurements. Because 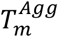 is likely to be a good measure of protein propensity to aggregate, and thus cause misfolding toxicity, we analyzed next its correlation with protein abundance and evolutionary rates.

To specifically analyze the potential effects of protein aggregation on protein evolution we used the 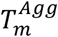 data for ~1500 *E. coli* proteins based on measurements performed in cells and natural cellular lysates (Mateus, et al. 2018). Interestingly, we found that in both datasets 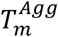 significantly correlated with protein abundances (**Figure 2A**, light grey; **Supplementary Table 2**; Spearman’s *r* = 0.20, *p* < 10^−14^, for cells; *r* = 0.21, *p* < 10^−15^, for cell lysates). However, we observed no significant correlations between 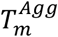 and evolutionary rates (**Figure 2A**, red). 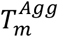 were also independently measured in two different *H. sapiens* cell lines: HeLa cells (Becher, et al. 2018), where the 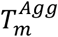 for ~4000 proteins were obtained for intact cells in different cell-cycle stages (G1/S transition and mitosis), and K562 chronic myeloid leukemia cells (Savitski, et al. 2014), where the 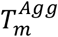 for ~2000 proteins were obtained for intact cells and cellular lysates. Analyzing these measurements, we found that across all human datasets 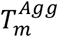 also positively correlated with protein abundances (**Figure 2B**, light grey; Spearman’s *r* = 0.20, *p* < 10^−38^; *r* = 0.19, *p* < 10^−32^; *r* = 0.29, *p* < 10^−36^; and *r* = 0.16, *p* < 10^−11^). For the K562 datasets, 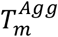 values were also negatively correlated with protein evolutionary rates (**Figure 2B**, red; Spearman’s *r* = −0.14, *p* < 10^−9^, for both cells and lysate). Finally, 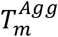 data were also obtained for ~800 proteins in Arabidopsis (Volkening, et al. 2019). In this case, we did not find any correlation of 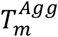 with protein abundances (**Figure 2C**, light grey) and the correlation with evolutionary rates was significant, but positive (**Figure 2C**, red; Spearman’s *r* = 0.18, *p* < 10^−6^).

**Figure 2.**
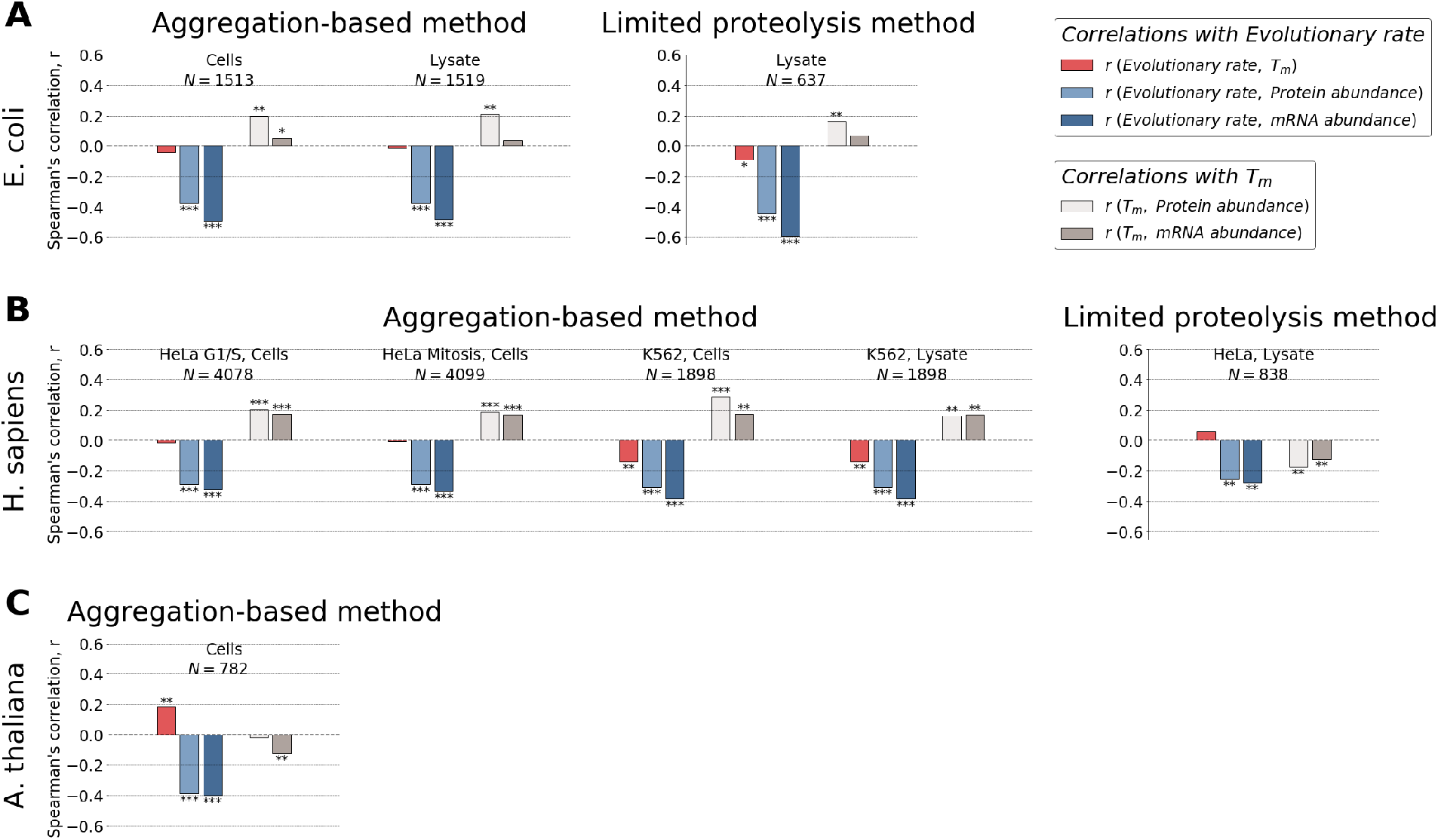
Correlations between genome-wide melting temperatures, protein abundances, mRNA abundances, and protein evolutionary rates. **A**, *E. coli*, **B**, *H. sapiens* and **C**, *A. thaliana.* Bar plots show the values of the Spearman’s correlation coefficients between evolutionary rates and melting temperatures (red), between evolutionary rates and protein abundances (light blue), between evolutionary rates and mRNA abundances (dark blue), between melting temperatures and protein abundances (light grey), and between melting temperatures and mRNA abundances (dark grey). Different methodologies used to measure *T*_*m*_ and different sample types are indicated above figure panels for *E. coli* and *H. sapiens*. Numbers of proteins in the analyzed datasets are shown; in each dataset we kept only proteins for which all four parameters are known. Asterisks above and below bars indicate significance level: * for p-value <0.05, ** for p-value <0.001, *** for p-value <10^−25^.

What fraction of the molecular clock variance is explained by the observed correlations with 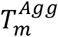? According to the MAH, avoiding cytotoxicity is a major driver of the variability in protein evolutionary rates. However, our analyses demonstrate that the anticorrelation between 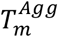 and evolutionary rates (**Figure 2**, red) is significant only in two (out of seven) datasets, and even in these two 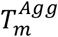 explain only ~2% of the variance in evolutionary rates across proteins. For comparison, mRNA abundance explains about an order of magnitude higher fraction of the evolutionary rate variance (**Figure 2**, dark blue), i.e. ~15% for the same subset of proteins. As expected due to weak correlations between 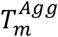 and evolutionary rates, we found that the anticorrelations between mRNA abundance and evolutionary rate do not substantially decrease after controlling for 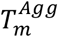 (for example, Spearman’s *r*_*Ev.Rate-mRNA*_ = −0.38 and corresponding Spearman’s partial 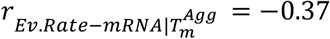 for *H. sapiens* K562 cells, the dataset with the strongest effects of 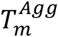).

To put the observed correlations with 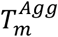 into perspective, we note that multiple other protein properties, such as the fraction of charged amino acids (Plata, et al. 2010), protein solubility (Plata, et al. 2010), surface stickiness (Levy, et al. 2012), and the number of protein-protein interaction partners (Yang, et al. 2012), have been shown to correlate with protein abundance. Changes in these properties with protein abundance detected in *E. coli* (Plata, et al. 2010; Levy, et al. 2012), *S. cerevisiae* (Levy, et al. 2012; Yang, et al. 2012) and *H. sapiens* (Levy, et al. 2012), likely help to alleviate deleterious effects of non-functional interactions and binding. For example, protein surface non-adhesiveness or the fraction of charged amino acids correlate positively, and with similar strength as 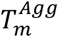, with protein abundances (**Figure 3**, light grey), and negatively with evolutionary rates (**Figure 3**, red, **Supplementary Table 3**, **Supplementary Table 4**). However, the ability of all these protein characteristics to explain the variability of evolutionary rates is modest compared to that of mRNA abundance (**Figure 3**, dark blue). Notably, protein surface non-adhesiveness, the fraction of charged amino acids and effects quantified by 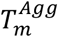 are likely to represent complementary sources of constraints, since they do not correlate strongly with each other (**Supplementary Table 5**).

**Figure 3.**
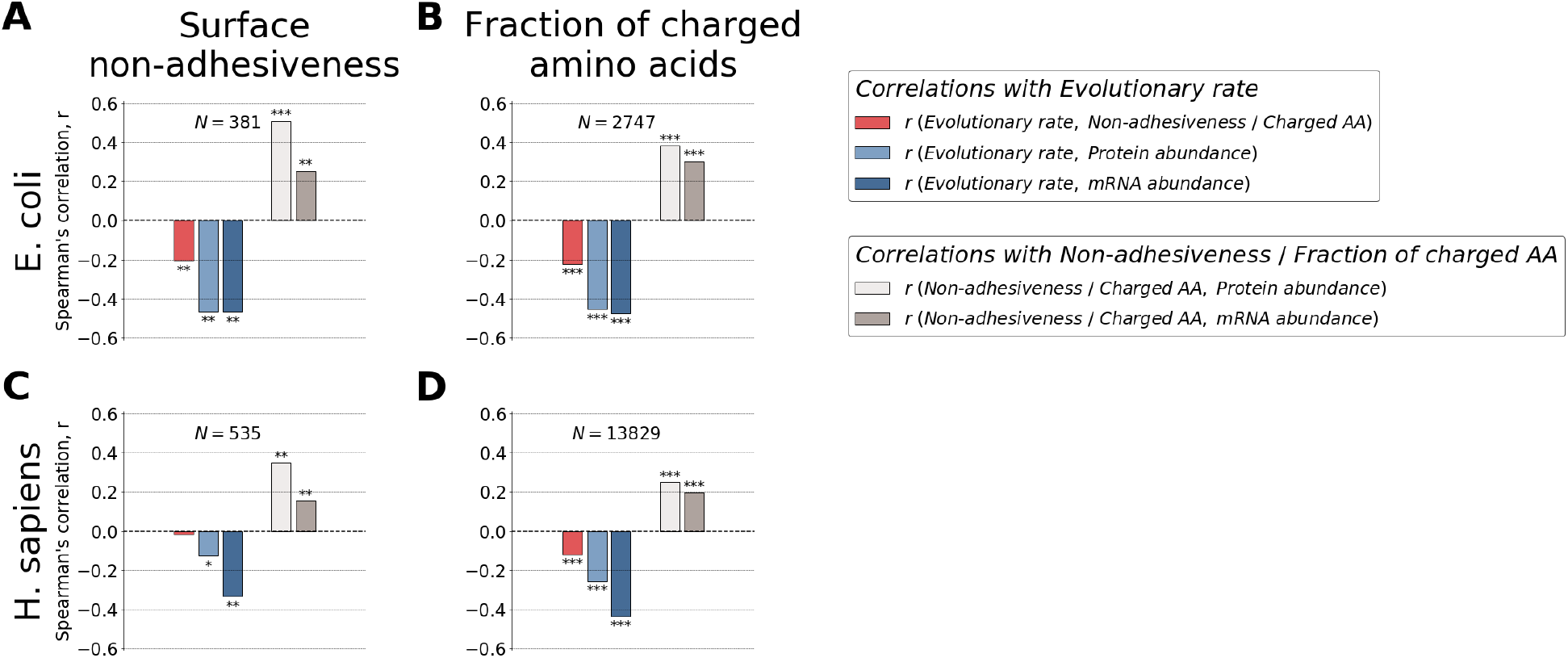
Correlations between protein surface non-adhesiveness, the fraction of charged amino acids, protein abundances, mRNA abundances, and protein evolutionary rates. **A, B**, *E. coli*, **C, D**, *H. sapiens*. Bar plots in **A** and **C** show the values of Spearman’s correlation coefficients between evolutionary rates and protein surface non-adhesiveness (red), between protein surface non-adhesiveness and protein abundance (light grey), and between protein surface non-adhesiveness and mRNA abundance (dark grey). Bar plots in **B** and **D** show the values of Spearman’s correlation coefficients between evolutionary rates and the fraction of charged amino acids (red), between the fraction of charged amino acids and protein abundance (light grey), and between the fraction of charged amino acids and mRNA abundance (dark grey). In all panels the values of Spearman’s correlation coefficients between evolutionary rates and protein abundances (light blue), and between evolutionary rates and mRNA abundances (dark blue) are also shown. Numbers of proteins in the analyzed datasets are indicated, in each dataset we kept only proteins for which all four parameters are known. Asterisks above and below bars represent significance level: * for p-value < 0.05, ** for p-value < 0.001, *** for p-value < 10^−25^.

## Discussion

In this work we continued testing the MAH using various empirical datasets, and the presented results agree with and extend our previous conclusions (Plata, et al. 2010; Plata and Vitkup 2018). The original MAH hypothesis was motivated, at least in part, by several studies showing that protein evolutionary rate was only weakly correlated with the fitness effect arising from complete deletion of the corresponding gene (Hurst and Smith 1999; Hirsh and Fraser 2001; Pal, et al. 2003). However, it is important not to conflate the effects associated with complete gene deletions and the level of overall protein sequence constraints which directly affect the rate of molecular clock. As discussed previously (Cherry 2010; Zhang and Yang 2015), protein sequence constraints arise due to selection against small fitness effect of single mutations, and the distribution of these small fitness effects usually does not reflect fitness loss associated with null mutations. Analogous observations have been made in different contexts. For example, although close yeast duplicates provide good buffering for complete knockouts of one homolog, individual amino acid mutations in close duplicates are actually more deleterious compared to mutations in genes without close homologs (Plata and Vitkup 2014).

A key question that the MAH was supposed to resolve is the nature of increased protein sequence constraints of highly abundant proteins. The original MAH (Drummond, et al. 2005; Drummond and Wilke 2008) and its multiple extensions (Yang, et al. 2010; Serohijos, et al. 2012) proposed that these constraints primarily originate from increased stability of highly expressed proteins. But, based either on direct measurements of protein stability (Δ*G*) or its various proxies (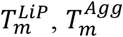, see **Supplementary Table 6**), we do not find support of these predictions. Proteins clearly need to be stable to perform their molecular and biological functions, and maintaining protein stability does constrain sequence evolution (Dill and Bromberg 2012). Multiple deep mutational scanning experiments demonstrate that fitness effects of substitutions correlate with their ΔΔ*G*, i.e. destabilizing mutations tend to be more deleterious (Jacquier, et al. 2013; Firnberg, et al. 2014; Sarkisyan, et al. 2016). Nevertheless, overall stability constraints are similar for different proteins, and differences in protein stability do not seem to play a major role in explaining the *variability* of evolutionary rates across cellular proteins. There is no paradox here, and this conclusion is in fact consistent with multiple empirical and biophysical data beyond the *T*_*m*_ and Δ*G* measurements. For example, it was demonstrated that the strength of the correlation between evolutionary rates and mRNA abundances is similar for sites with different contributions to protein stability, such as surface sites and sites in protein cores (Yang, et al. 2012). Moreover, increasing protein stability beyond a certain threshold is not evolutionary advantageous, and may be generally detrimental to fitness, as demonstrated by multiple examples of stability-activity tradeoffs in proteins (Wang, et al. 2002; Tokuriki, et al. 2008; Knies, et al. 2017). If effects associated with misfolding become harmful, for example due to significantly increased burden of transcriptional (Goldsmith and Tawfik 2009) or translational (Bratulic, et al. 2015) errors, proteins can be quickly stabilized by fixation of several mutations (Goldsmith and Tawfik 2009; Bratulic, et al. 2015) without substantial further constraints on the corresponding protein sequence.

Another important question related to the MAH is understanding the primary origin of protein expression costs. In the original MAH hypothesis this cost was proposed to arise from protein misfolding induced by translational errors (Drummond, et al. 2005; Drummond and Wilke 2008), and this cost was later extended to error free misfolding (Yang, et al. 2010). We note that several experimental studies did not find substantial costs associated with protein misfolding (Plata, et al. 2010; Kafri, et al. 2016). By overexpressing pairs of close yeast duplicates, evolving at different rates, a recent study by Biesiadecka *et al.* (Biesiadecka, et al. 2020) also did not find any evidence of substantial contribution of costs associated with translation-induced misfolding. Although the study by Geiler-Samerotte *et al.* (Geiler-Samerotte, et al. 2011) is often viewed as supporting a major role of the MAH (Zhang and Yang 2015), the results reported in that study also suggest a smaller cost of protein misfolding compared to the costs associated with protein production. Geiler-Samerotte *et al.* showed that the gratuitous overexpression of a protein with multiple destabilizing substitutions leads to deleterious fitness effects which are ~3 times higher compared to the expression cost of the wild-type protein. However, mistranslation errors are present in only ~15% of proteins (Drummond and Wilke 2008; Yang, et al. 2010), and even a smaller fraction of proteins contains multiple translation-induced mutations or mutations with substantial destabilizing effects (for example, only ~20% of mutations have ΔΔ*G* < −2 kcal/mol) (Nisthal, et al. 2019). Therefore, the overall cost of translational-induced misfolding (per protein) is substantially smaller than the cost of protein production. The same conclusion can be extended to error-free misfolding based on the estimate that it contributes only 5-20% extra misfolding events compared to misfolding arising from translational errors (Yang, et al. 2010).

Interestingly, across the *E. coli* and *H. sapiens* datasets we analyzed, 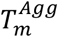 correlates better with protein abundances, while evolutionary rates correlate more strongly with mRNA abundances (**Figure 2**). This result again suggests that different biological mechanisms may be driving sequence constraints related to protein aggregation and those responsible for the substantial variance of protein evolutionary rates. Direct experimental measurements demonstrated that deleterious mutations reduce fitness primarily due to changes in protein function, rather than due to destabilization-induced changes in protein abundance (Firnberg, et al. 2014). Analyzing long-term protein evolution, we also recently showed that functional optimality, i.e. the conservation of protein sequence and 3-dimensional structure necessary for efficient protein function, is a substantially stronger evolutionary constraint than the requirement to simply maintain folded protein stability (Konate, et al. 2019). Recently, the fraction of mutations leading to deleterious effects through all possible non-functional mechanisms, called collateral effects, was estimated to be ~40% for the TEM-1 protein in *E. coli* (Mehlhoff, et al. 2020). This result also suggests that collateral effects are unlikely to dominate protein evolutionary constraints, at least for the vast majority of bacterial proteins. Based on the ratio of amino acid changing to synonymous mutations in bacteria, *K*_a_/*K*_s_~0.05-0.1 (Koonin 2012) the fraction of all bacterial amino acid changing mutations that are rejected in evolution is ~0.9-0.95. Therefore, functional effects play a larger role in purifying selection even under the assumption that collateral mechanisms dominate functional mechanisms for all mutations with detected collateral effects. And this assumption is quite unlikely as protein sites of collateral mutations substantially overlap with functionally-sensitive sites, and collateral effects are often smaller in magnitude compared to functional effects (Stiffler, et al. 2015; Mehlhoff, et al. 2020). All these results suggest that the diversity of protein evolutionary rates may be more related to functional effects of mutations rather than effects associated with protein stability. Computational models (Cherry 2010; Gout, et al. 2010), also demonstrated the plausibility that by minimizing costs associated with protein production, functional optimization may be responsible for higher level of sequence constraints of abundant cellular proteins.

Finally, although the feasibility of a dominant MAH contribution was suggested by computational simulations (Drummond and Wilke 2008; Yang, et al. 2010; Serohijos, et al. 2012), biology is an empirical science, and the fidelity of proposed hypotheses should be ultimately determined by their agreement, or lack thereof, with available experimental data. An obvious weakness of aforementioned simulations is that they only considered effects associated with protein stability and interactions. Thus, they demonstrated the MAH feasibility but could not evaluate the relative importance of other biological and functional effects. We also note that it is often possible to invoke sophisticated noise and error models to explain the absence of expected observations (Razban 2019). Nevertheless, based on the preponderance of available evidence, the MAH is unlikely to play any major role in explaining a strong negative correlation between protein abundance and evolutionary rate. Major roles are also unlikely for several other effects, such as increased protein solubility or avoidance of non-functional interactions (Plata, et al. 2010; Yang, et al. 2012). Most importantly, the search for the main factors contributing to the substantial variability of evolutionary rates across proteins must continue.

## Materials and Methods

The protein stability data were obtained from the ProTherm database (Kumar, et al. 2006), which have been recently moved to the ProtaBank (Wang, et al. 2019) and are available at https://github.com/protabit/protherm-conversion. We considered unfolding Gibbs free energies, Δ*G*, for wild type proteins measured at pH values between 4 and 9, and for temperatures between 10 and 50 degrees C. Proteins in all oligomeric states and with any (un)folding dynamics were used in Figure 1. Only monomers that exhibit 2-state reversible (un)folding were used in Supplementary Figure 1. For each protein with more than one available measurement the values were averaged over different experimental conditions and measurements performed by different research groups. Raw Δ*G* values for *E. coli* and *H. sapiens* extracted from ProTherm are available in **Supplementary Table 7**.

We used the rate of non-synonymous substitutions, *K*_a_, as a measure of protein evolutionary rate. *K*_a_ values for *Escherichia coli*, *Homo sapiens* and *Arabidopsis thaliana* were calculated using the PAML package (Yang 1997) relative to *Salmonella enterica*, *Mus musculus* and *Brassica oleracea*. Orthologues were identified as bidirectional best hits in pairwise local alignments calculated with Usearch (Edgar 2010). We considered for analysis only protein pairs for which corresponding alignments covered at least 70% of the shortest protein length.

Protein abundance data for all species were obtained from the whole-organism integrated datasets available in the PaxDB v.4 database (Wang, et al. 2015). mRNA abundances data for *E. coli* were obtained from (McClure, et al. 2013). mRNA abundances from the brain frontal cortex (Mele, et al. 2015) was used for *H. sapiens*, since it was demonstrated that mRNA expression in this tissue has the highest correlation with protein evolutionary rates (Drummond and Wilke 2008). mRNA abundances from the germinating seed was used for *A. thaliana* (Klepikova, et al. 2016).

Protein surface non-adhesiveness was obtained from (Levy, et al. 2012). Specifically, this measure equals the negative sum of amino acid stickiness scores (Levy, et al. 2012) across sequence sites located on protein surfaces based on protein 3-dimentional structures (Levy 2010).

The data used for analyses in the manuscript are available in **Supplementary Tables 8-10**.

## Acknowledgements

We would like to thank Drs. Purushottam Dixit and Balázs Papp for helpful scientific discussions. This work was supported by the National Institute of General Medical Sciences grant R35GM131884 to Dennis Vitkup.

---

**Supplementary Figure 1. Correlations between experimentally measured Δ*G* values, protein abundances, mRNA abundances, and protein evolutionary rates for monomers with 2-state reversible (un)folding**

**A.** *E. coli* (n=24) and **B.** *H. sapiens* (n=38). The correlations between evolutionary rates and unfolding Gibbs free energies, Δ*G*, are shown in the first figure column (red). The correlations between protein evolutionary rates and mRNA abundances are shown in the second column (blue). The correlations between Δ*G* and protein abundances are shown in the third column (light grey), and the correlations between Δ*G* and mRNA abundances are shown in the fourth column (dark grey). Solid lines represent the least square regressions fitted to the data. Spearman’s correlation coefficients and corresponding p-values are shown, significant correlations are highlighted in bold. Only Δ*G* values for monomers which exhibit 2-state reversible (un)folding are included in this analysis.

**Supplementary Table S1**

Correlations between experimentally measured ΔG values, protein abundances, mRNA abundances, and protein evolutionary rates

**Supplementary Table S2**

Correlations between genome-wide melting temperatures, protein abundances, mRNA abundances, and protein evolutionary rates

**Supplementary Table S3**

Correlations between protein surface non-adhesiveness, protein abundances, mRNA abundances, and protein evolutionary rates

**Supplementary Table S4**

Correlations between fractions of charged amino acids, protein abundances, mRNA abundances, and protein evolutionary rates

**Supplementary Table S5**

Correlations between protein surface non-adhesiveness, fractions of charged amino acids, and melting temperatures

**Supplementary Table S6**

Correlations between different proxies of protein stability (ΔG, 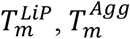)

**Supplementary Table S7**

ΔG values for *E. coli* and *H. sapiens* extracted from ProTherm

**Supplementary Table S8**

Data on evolutionary rates, mRNA abundances, protein abundancies, ΔG, melting temperatures, surface non-adhesiveness, and fractions of charged amino acids for *E. coli*.

**Supplementary Table S9**

Data on evolutionary rates, mRNA abundances, protein abundancies, ΔG, melting temperatures, surface non-adhesiveness, and fractions of charged amino acids for *H. sapiens*.

**Supplementary Table S10**

Data on evolutionary rates, mRNA abundances, protein abundancies, melting temperatures, and fractions of charged amino acids for *A. thaliana*.

