## Supplementary figures and images for "The relationship between misfolding avoidance hypothesis and protein evolutionary rates in the light of empirical evidence"

### Supplementary figure 1

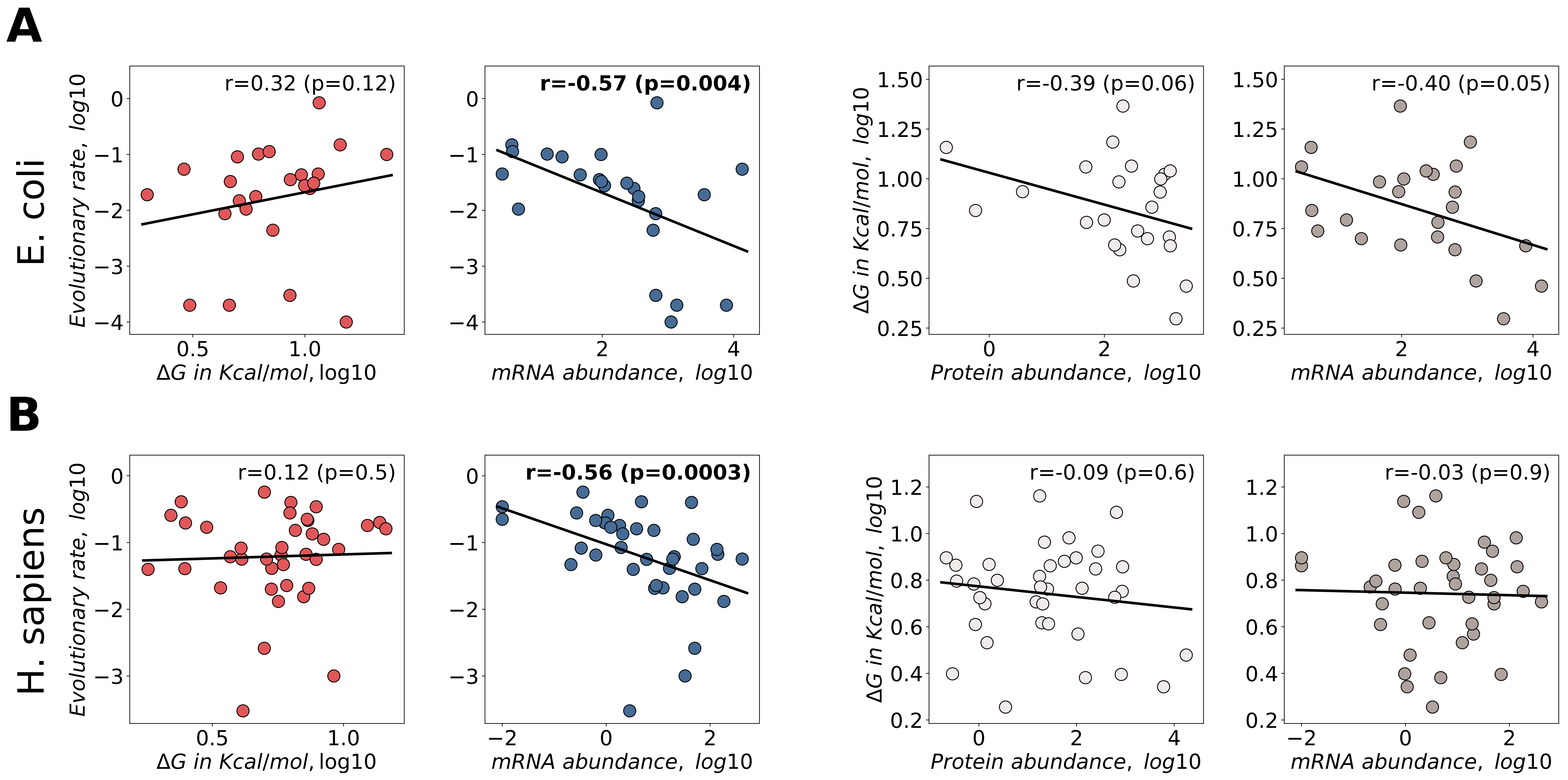
